# The Impact of Preprints on the Citations of Journal Articles Related to COVID-19

**DOI:** 10.1101/2024.07.21.604465

**Authors:** Hiroyuki Tsunoda, Yuan Sun, Masaki Nishizawa, Xiaomin Liu, Kou Amano, Rie Kominami

## Abstract

To investigate the impact of preprints on the citation counts of COVID-19-related papers, this study compares the number of citations received by drafts initially distributed as preprints and later published in journals with those received by papers directly submitted to journals. The difference in the median number of citations between COVID-19 preprint-distributed papers and COVID-19 directly submitted papers published in 184 journals was tested using the Mann-Whitney U test. The results showed that 129 journals had a statistically significant higher median citation count for COVID-19 preprint-distributed papers compared to directly submitted papers, with a p-value of less than 0.05. In contrast, no journals had a statistically significant higher median citation count for COVID-19 directly submitted papers. This indicates that 70.11% of the journals that published preprint-distributed papers experienced a significant increase in citations. We also identified that among the 184 journals, 13 journals garnered a substantial number of citations. Among the 74,037 COVID-19 papers, preprint-distributed papers (9,028) accounted for only 12.19%. However, among the 2,015,997 citations received by COVID-19 papers, preprint-distributed papers garnered 542,715 citations, representing a substantial 26.92%. These results suggest that distributing preprints prior to formal publication may help COVID-19 research reach a wider audience, potentially leading to increased readership and citations.

## 1 Introduction

Preprints are openly accessible scientific papers distributed from an archive before they are submitted to a journal and undergo peer review. The value and importance of preprints have increased since they contributed to the public health emergency of the COVID-19 pandemic, and their role as a new means of academic information dissemination is now anticipated [1, 2, 3]. However, although preprints are increasing rapidly, they remain confined to certain regions, fields, and subjects, and have not yet spread throughout the academic community [4]. One reason authors are hesitant to submit to preprints is the concern about publicly sharing research that has not undergone the peer review process. Moreover, many researchers are not even aware of preprints [5]. Some researchers have raised concerns about the quality of preprints since they have not undergone peer review. Recent studies have shown that the content of preprints generally matches that of the papers once they are submitted to journals, peer-reviewed, and published [6, 7, 8, 9]. Additionally, many preprints are published in high-impact journals included in Clarivate Analytics’ Web of Science Core Collection, indicating that preprints are not uniformly of low quality [10]. Studies on the peer review period have shown that research papers initially distributed as preprints tend to undergo a shorter peer review process when submitted to journals, leading to faster publication [11, 12]. Preprints indicate that papers related to COVID-19 are published to journals at a higher frequency than those unrelated to COVID-19 [13, 14] and undergo peer review more expeditiously [15, 16]. Studies on dissemination have shown that preprints are extensively covered by the media [17, 18] and are widely disseminated on social media [19]. The wide dissemination of preprints may be a factor in the subsequent citation of the papers after peer review. Furthermore, in the medical field, authors tend to submit COVID-19-related papers, initially distributed as preprints, to journals with high impact factors [20], and these papers receive numerous citations. For example, as of June 2024, in citations of articles indexed in PubMed and assigned PubMed IDs, the paper by Cummings et al. first opened on medRxiv on April 20, 2020, and later published in The Lancet on May 19, 2020, received 1,165 citations [21,22]. The paper by Guan et al. first opened on medRxiv on February 9, 2020, and published in the New England Journal of Medicine on April 30, 2020, received 14,683 citations [23,24]. The manuscript by Wang et al., first opened on medRxiv on March 6, 2020, was revised with Pan as the first author and a new title for journal submission, and published in JAMA on April 10, 2020, received 853 citations [25,26]. Whether the high number of citations is due to individual preprints or the characteristics of preprints themselves remains unclear. The purpose of this study is to compare the number of citations received by drafts initially distributed as preprints that were later published in journals with those received by papers directly submitted to journals, in order to elucidate the impact of preprints on the citation of papers.

## 2 Data and Method

### 2.1 Preprints

A preprint is a draft that is made publicly available in an archive before being submitted to a journal. Preprints have not yet undergone peer review. In this study, we will use preprints from two archives, medRxiv and bioRxiv, which focus on medical and biological topics, to investigate papers related to COVID-19. On January 30, 2020, the World Health Organization (WHO) declared the novel coronavirus infection a “Public Health Emergency of International Concern (PHEIC).” Subsequently, considering the global spread and severity of the infection, the WHO declared it a pandemic on March 11, 2020.

### 2.2 Journal Articles Related to COVID-19

PubMed is a free resource supporting the search and retrieval of biomedical and life sciences literature with the aim of improving global and individual health. The PubMed contains over 37 million citations and abstracts of biomedical literature. Available to the public online since 1996, PubMed was developed and maintained by the National Center for Biotechnology Information (NCBI) at the U.S. National Library of Medicine (NLM), which is part of the National Institutes of Health (NIH) [27]. In this study, to comprehensively collect papers related to COVID-19, we gathered papers published on PubMed. For collecting the papers, we used the Entrez Programming Utilities (E-utilities), a publicly accessible Application Programming Interface (API) for the NCBI Entrez system, and the program was written in Python. The determination of the subjects of the papers was made using the Medical Subject Headings (MeSH) thesaurus created by the National Library of Medicine. Papers recorded with “COVID-19” in MeSH were defined as COVID-19 related papers.

### 2.3 Collection Period and Types of Documents

Since papers on topics related to COVID-19 had already been published in journals as early as 2019, the paper collection period was set from 2019 to 2023. Journals publish articles of various types. Here, we defined the document types as journal articles and reviews to measure impact using citation counts.

### 2.4 Citation Counts

In citation analysis, the citation counts often used are the “Times Cited” produced by Clarivate and provided by WoS. Measuring impact requires investigating a large number of papers, but Clarivate imposes restrictions on downloading papers programmatically. Therefore, in this study, we utilized PubMed, which allows citation downloads via API. When a paper published on PubMed is cited by another paper on PubMed, the citing paper is displayed in the “Cited by” section. The citation count of a paper is determined by counting the number of citing papers.

### 2.5 Analysis and Examination of Citation Counts

There are several methods for assigning citation counts to a collection of papers gathered by media such as journals or by subject. The total citation count of the papers that constitute the elements of the collection, the average obtained by dividing the total citation count by the number of elements, and the median citation count of the papers are potential candidates for representing the citation count of the collection. The total citation count is not adopted because the number of elements in the collections varies significantly. The distribution of citation counts in the paper collections is not normal; there are many papers with low citation counts and few papers with high citation counts, indicating a non-normal distribution. The average citation count is strongly influenced by a few highly-cited papers. On the other hand, the median is not affected by these outliers and represents the relative position of the entire collection. Therefore, the median is adopted. For testing the difference in citation counts, we employ the Mann-Whitney U test, a non-parametric test used for collections with significantly different sizes and non-normal distributions. A p-value of less than 0.05 is considered statistically significant.

### 2.6 Journal Selection

First, preprints that had been submitted to journals, accepted, and assigned a publisher’s DOI were identified. Next, among the journals that published papers initially disseminated as preprints, those indexed in PubMed were identified. Finally, from these, the journals with a high number of papers initially disseminated as preprints were selected for the investigation.

## 3 Result

### 3.1 Status of the Paper

As for the journals selected for investigation, 184 journals that published at least 10 papers initially disseminated as preprints were identified from PubMed. Metadata for 952,973 papers published by these journals between 2019 and 2023 were collected from PubMed. The total number of citations these papers received from other papers indexed in PubMed was 9,393,014

### 3.2 COVID-19 Papers and Non-COVID-19 Papers

The median number of citations for COVID-19 and non-COVID-19 papers published in 184 journals was tested using the Mann-Whitney U test. Among the COVID-19 papers, 155 journals showed a significant increase in citations with a p-value of less than 0.05. Conversely, for non-COVID-19 papers, only 2 journals showed a significant increase in citations. This means that 84.24% of the journals that published COVID-19 papers experienced an increase in citations. This situation is shown in Table 1.

**Table 1.** COVID-19 Papers and Non-COVID-19 Journals.

| Difference | Significant | (%) | Non-Significant | (%) | Journals |
| --- | --- | --- | --- | --- | --- |
| Positive | 155 | 84.24 | 8 | 4.35 | 163 |
| Zero | 8 | 4.35 | 6 | 3.26 | 14 |
| Negative | 2 | 1.09 | 5 | 2.72 | 7 |
|  | 165 |  | 19 |  | 184 |

Of the 952,973 papers, 74,037 were assigned the MeSH term COVID-19 (hereafter referred to as COVID-19 papers), with a total citation count of 2,015,997. On the other hand, 878,936 papers were not assigned this term (hereafter referred to as non-COVID-19 papers), with a total citation count of 7,377,017. Thus, 7.77% of the COVID-19 papers accounted for 21.46% of the citations. This situation is shown in Table 2.

**Table 2.** COVID-19 Papers and Non-COVID-19 Papers.

|  | Papers | (%) | Citations | (%) |
| --- | --- | --- | --- | --- |
| COVID-19 | 74,037 | 7.77 | 2,015,997 | 21.46 |
| Non-COVID-19 | 878,936 | 92.23 | 7,377,017 | 78.54 |
|  | 952,973 |  | 9,393,014 |  |

### 3.3 Papers initially disseminated as preprints and direct journal submission papers

The difference in the median number of citations between preprint-disseminated papers and direct-submitted papers published in 184 journals was tested using the Mann-Whitney U test. For preprint-disseminated papers, the p-value was less than 0.05, indicating that there were 161 journals with significantly higher citation counts. In contrast, there were only 3 journals with significantly higher citations for direct-submitted papers. This means that 87.50% of the journals that published preprint-disseminated papers experienced an increase in citations. This situation is shown in Table 3.

**Table 3.** Preprint-disseminated and direct-submitted journals.

| Difference | Significant | (%) | Non-Significant | (%) | Journals |
| --- | --- | --- | --- | --- | --- |
| Positive | 161 | 87.50 | 12 | 6.52 | 173 |
| Zero | 2 | 1.09 | 4 | 2.17 | 6 |
| Negative | 3 | 1.63 | 2 | 1.09 | 5 |
|  | 166 |  | 18 |  | 184 |

Out of 952,973 papers, the number of papers initially disseminated as preprints (hereafter referred to as preprint-disseminated papers) was 54,909, with a total of 1,293,049 citations. On the other hand, the number of papers submitted directly to journals (hereafter referred to as direct-submitted papers) was 898,064, with a total of 8,099,965 citations. The 5.76% of preprint-disseminated papers garnered 13.77% of the citations. This situation is shown in Table 4.

**Table 4.** Preprint-disseminated and direct-submitted papers.

|  | Papers | (%) | Citations | (%) |
| --- | --- | --- | --- | --- |
| Preprint-disseminated | 54,909 | 5.76 | 1,293,049 | 13.77 |
| direct-submitted | 898,064 | 94.24 | 8,099,965 | 86.23 |
|  | 952,973 |  | 9,393,014 |  |

### 3.4 Papers initially disseminated as preprints and direct journal submission papers related to COVID-19

The difference in the median number of citations between COVID-19 preprint distribution papers and COVID-19 directly submitted papers published in 184 journals was tested using the Mann-Whitney U test. For COVID-19 preprint distribution papers, the p-value was less than 0.05, indicating that there were 129 journals with significantly higher citation counts. In contrast, there were no COVID-19 directly submitted papers with significantly higher citations. This means that 70.11% of the journals that published preprint distribution papers experienced an increase in citations. This situation is shown in Table 5.

**Table 5.** COVID-19 Preprint-disseminated and direct-submitted papers.

| Difference | Significant | (%) | Non-Significant | (%) | Journals |
| --- | --- | --- | --- | --- | --- |
| Positive | 129 | 70.11 | 44 | 23.91 | 173 |
| Zero | 0 | 0 | 3 | 1.63 | 3 |
| Negative | 0 | 0 | 8 | 4.35 | 8 |
|  | 129 |  | 55 |  | 184 |

Out of 74,037 COVID-19 papers, the number of preprint distribution papers (referred to as COVID-19 preprint distribution papers) was 9,028, with a total of 542,715 citations. On the other hand, the number of papers directly submitted to journals (referred to as COVID-19 directly submitted papers) was 65,009, with a total of 1,473,282 citations. Thus, 12.19% of the preprint distribution papers accounted for 26.92% of the citations. This situation is shown in Table 6.

**Table 6.**
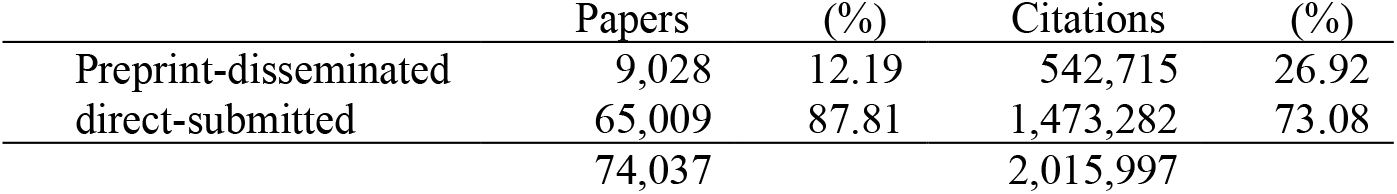
COVID-19 Preprint-disseminated and direct-submitted papers.

Table 7 shows the number of COVID-19 preprint-disseminated papers and direct-submitted papers in journals, the total number of citations, the median number of citations, the p-values, the significance, and the differences.

**Table 7.**
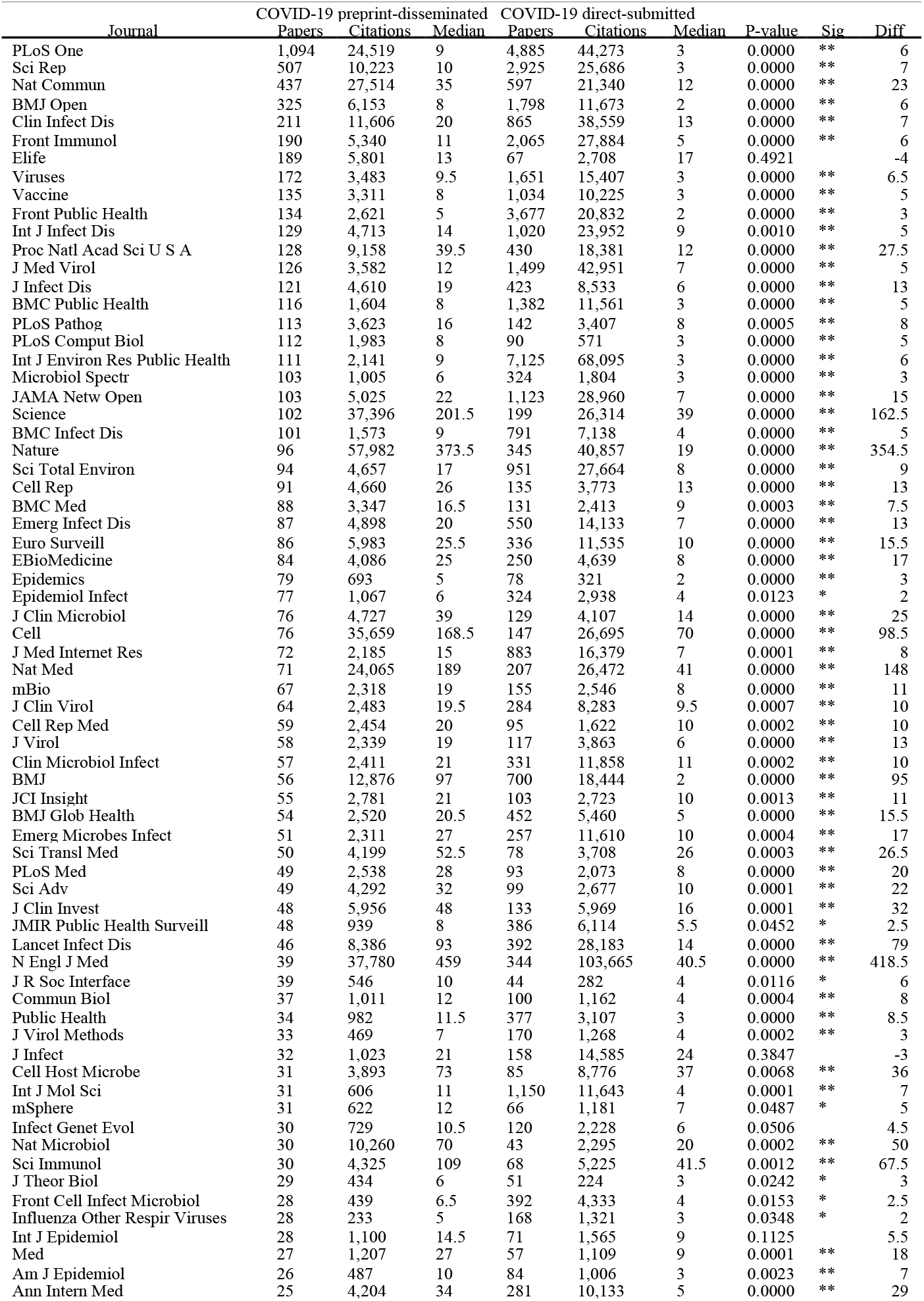
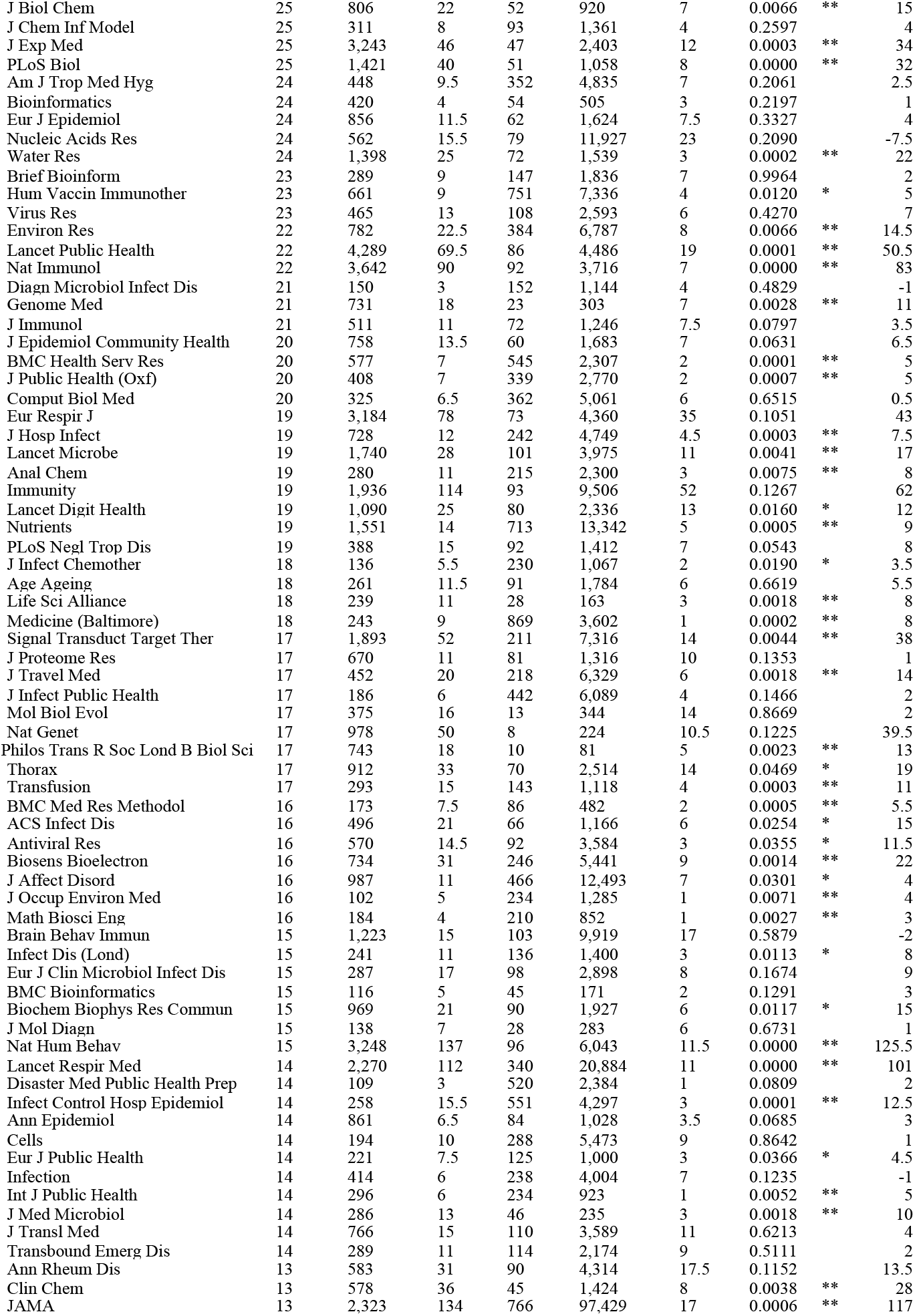
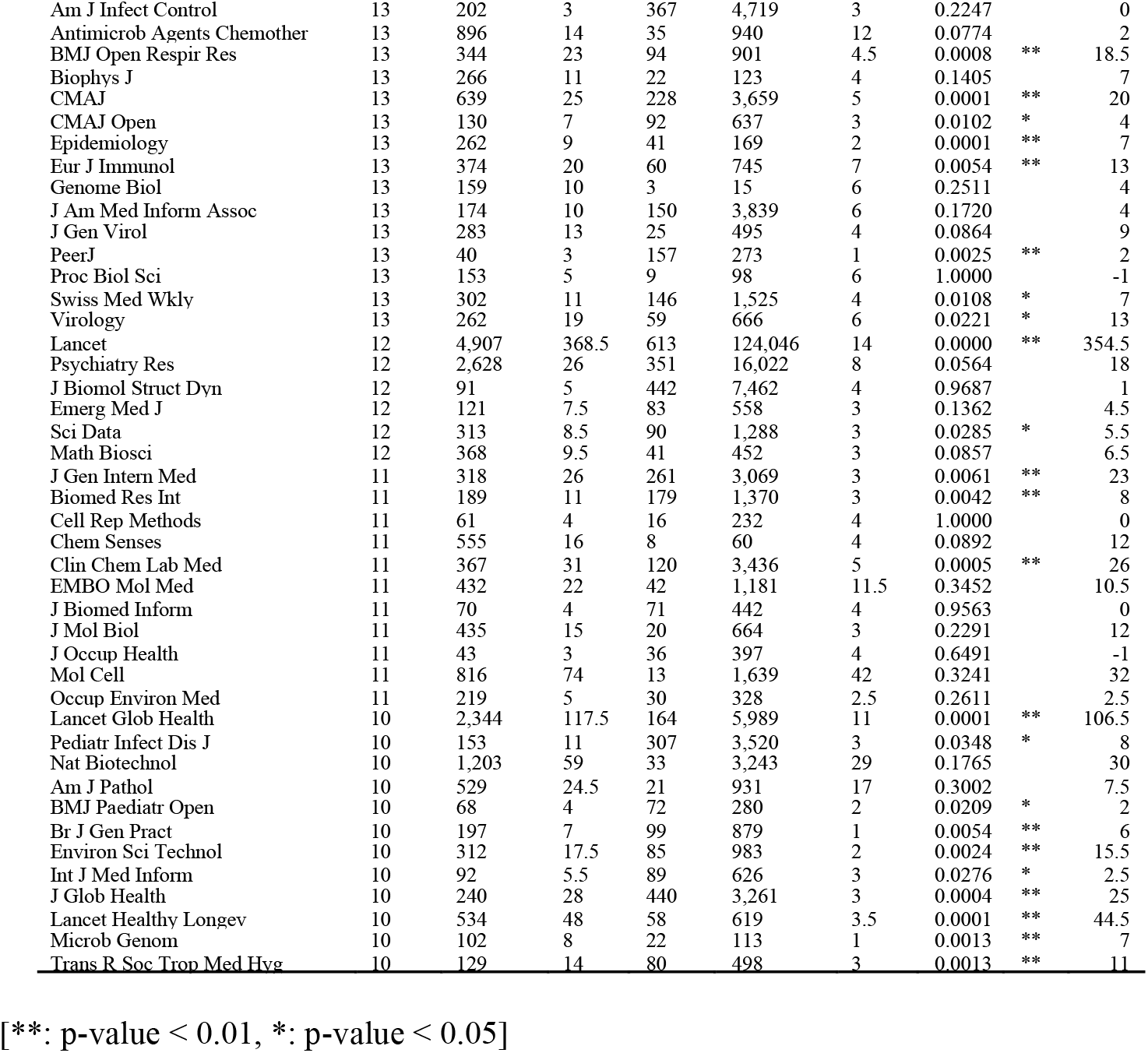
COVID-19 preprint-disseminated and direct-submitted papers.

## 4 Analysis and Discussion

### 4.1 Distribution of the median number of citations by journal

Figure 1 shows the distribution of the median number of citations for COVID-19 papers and non-COVID-19 papers, preprint-disseminated and direct-submitted papers, and COVID-19 preprint-disseminated and direct-submitted papers. Figure 1 shows that preprint-distributed papers have higher citation counts than COVID-19 papers, and that COVID-19 preprint-distributed papers tend to have even higher citation counts.

**Fig. 1.**
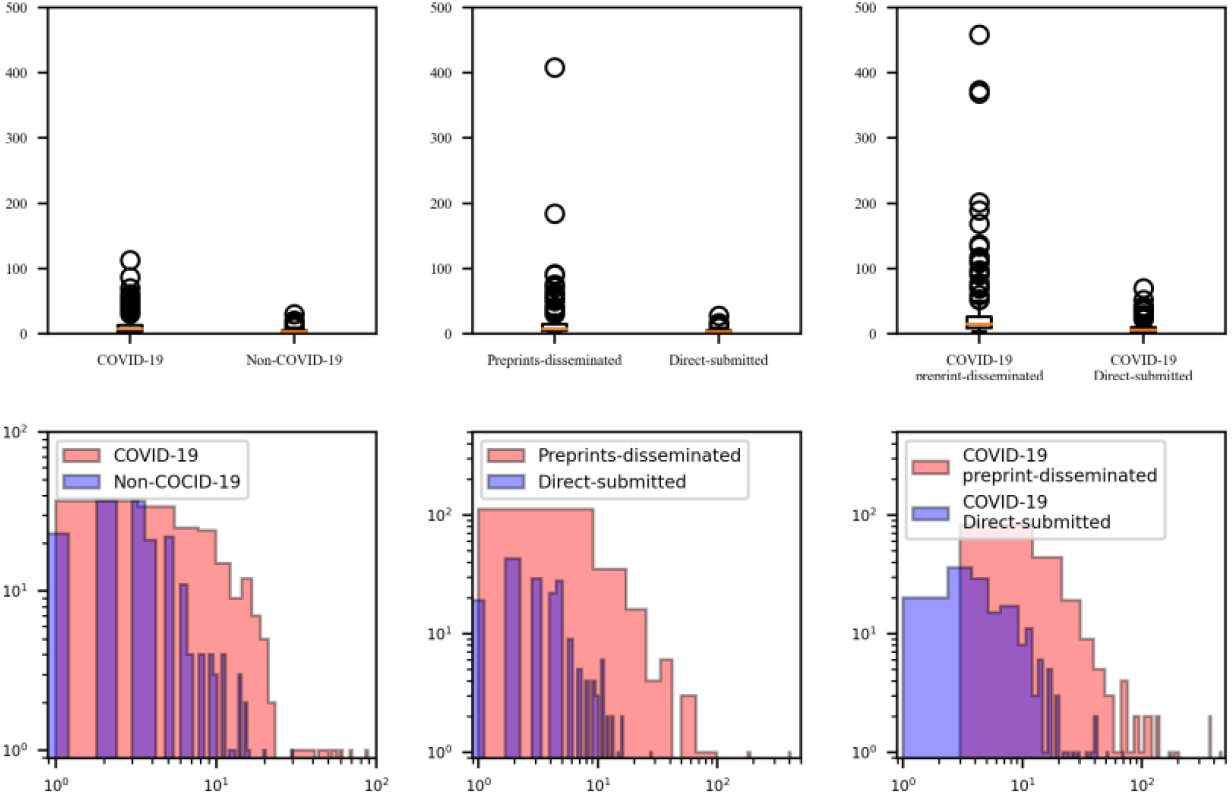
Distribution of the median number of citations.

### 4.2 Relative position of journals based on the distribution of median citation counts

In journals with a higher median citation count for preprint distribution papers compared to papers directly submitted to journals—i.e., journals with a positive difference—clustering with a citation threshold of 20 resulted in seven clusters. The first cluster consisted of a single journal, *The New England Journal of Medicine*. The second cluster included two journals, *The Lancet* and *Nature*. The third cluster comprised one journal, *Cell*. The fourth cluster included two journals, *Science* and *Nature Medicine*. The fifth cluster consisted of seven journals: *Nature Immunology, The Lancet Infectious Diseases, BMJ, The Lancet Global Health, The Lancet Respiratory Medicine, JAMA*, and *Nature Human Behaviour*. The seventh cluster included two journals, *Immunity* and *Science Immunology*. Finally, the sixth cluster included the remaining 158 journals. The 13 journals classified into clusters 1 through 5 were found to have garnered an especially large number of citations. This cluster is illustrated in the dendrogram shown in Figure 2.

**Fig. 2.**
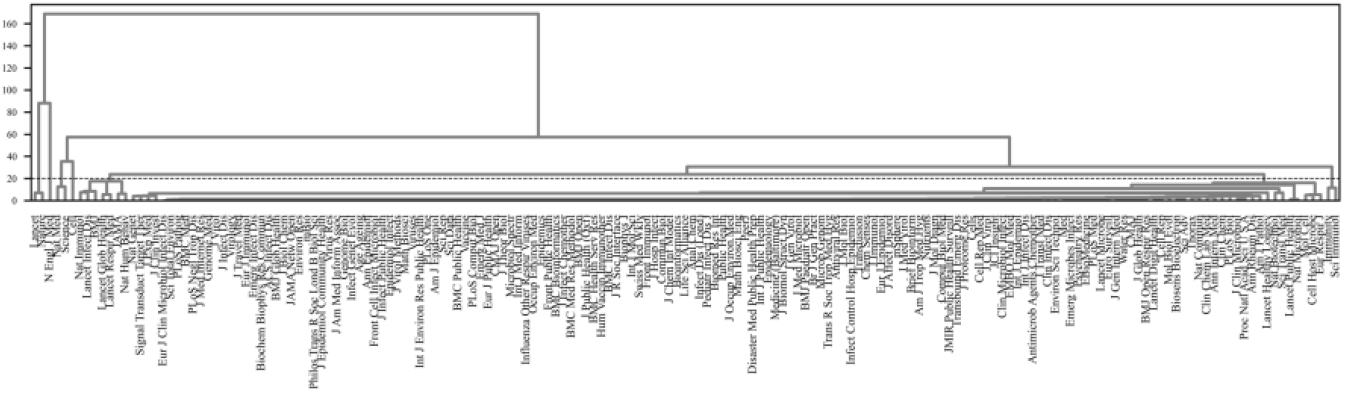
Positive median difference in citation counts of journal articles on COVID-19.

Figure 3 shows the diagram obtained by adding two clusters with zero and negative differences to the seven clusters with positive differences shown in Figure 2. Clearly, among the seven positive clusters, clusters 1 through 5 are located in the upper-left part of the figure 3, visually confirming that distributing the draft as a preprint significantly increased the number of citations.

**Fig. 3.**
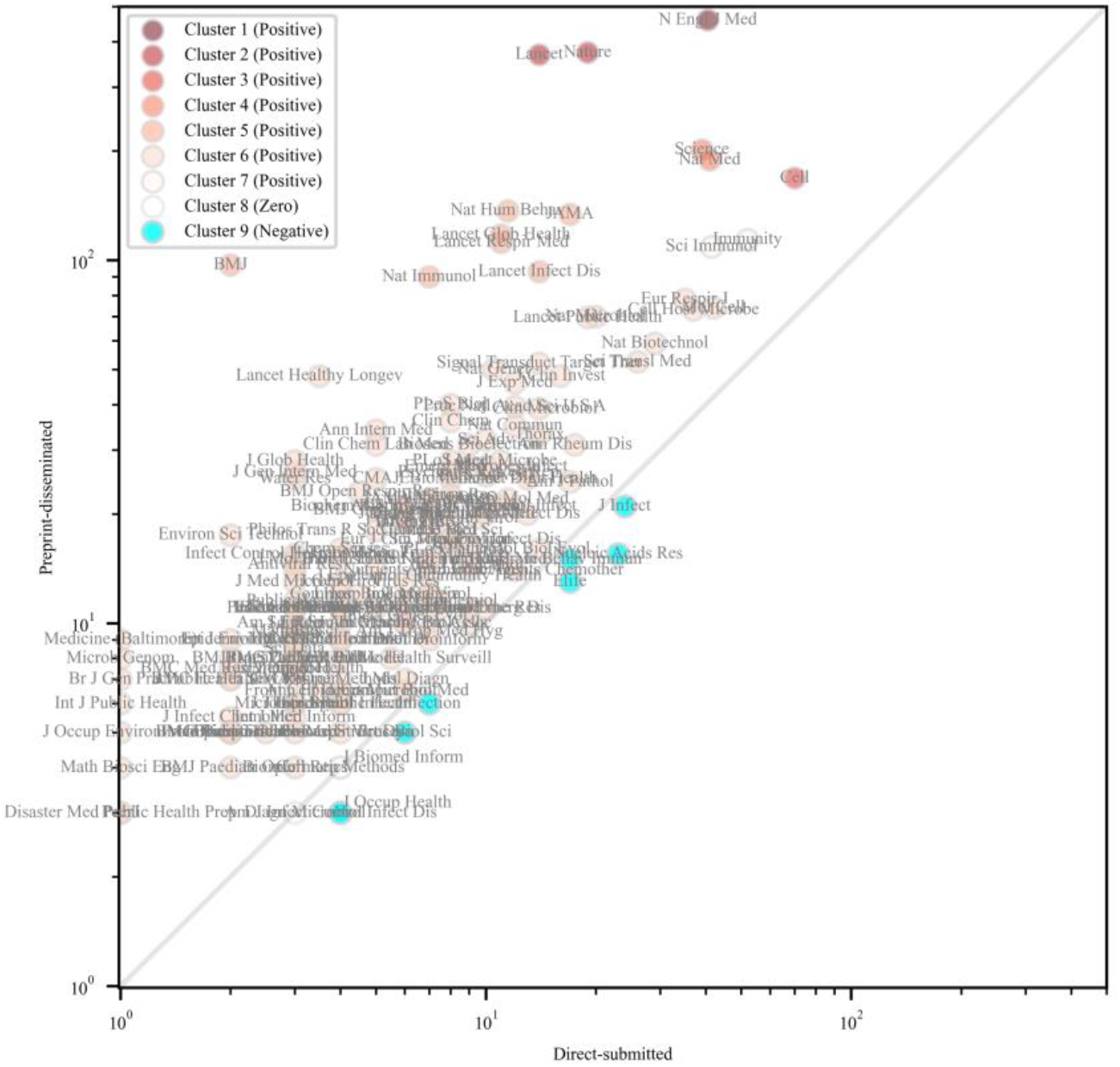
Median citation count of journal articles on COVID-19.

### 4.3 Impact of Preprints on the Citations of Journal Articles Related to COVID-19

In most journals, COVID-19 papers received more citations than non-COVID-19 papers. Furthermore, papers first distributed as preprints received more citations than those directly submitted to journals by the authors. It can be inferred that the prior dissemination of preprints allowed the research to reach a wider audience, thereby gaining more readers and citations.

## 5 Conclusion

In this study, we analyzed the number of times PubMed articles were cited by other PubMed articles to elucidate the impact of preprints on COVID-19 research. The results revealed that, among papers on COVID-19, preprint-distributed papers received significantly more citations than direct journal submission papers, as demonstrated by the analysis of the median difference in journal citation counts. This indicates that distributing preprints exerts a strong influence on the subsequent citation counts of papers published in journals. For novel topics like COVID-19, it is crucial for researchers to rapidly disseminate their findings. Therefore, researchers are encouraged to utilize preprints to share their research outcomes promptly.

## Acknowledgments

This work was supported by JSPS KAKENHI Grant Numbers JP19K12707, JP20K12569, JP22K12737, and ROIS NII Open Collaborative Research 2023(23FS01)

## Disclosure of Interests

The authors have no conflicts of interest directly relevant to the content of this article.

## Notes

### Competing Interest Statement

The authors have declared no competing interest.

